# Effects of six pyrimidine analogs on the growth of *Tetrahymena thermophila* and their implications in pyrimidine metabolism

**DOI:** 10.1101/2023.03.29.534814

**Authors:** Zander Harpel, Wei-Jen Chang, Jacob Circelli, Richard Chen, Ian Chang, Jason Rivera, Stephanie Wu, Zonhan Wei

**Affiliations:** Department of Biology, Hamilton College, Clinton, 13323, NY, USA; New Hartford Senior High School, New Hartford, 13413, NY, USA; Clinton Senior High School, Clinton, 13323, NY, USA; School of Mechanics and Engineering Science, Zhengzhou University, Zhengzhou, 450001, Henan, China

**Keywords:** Endocytosis, membrane trafficking, nucleoside transport, pyrimidine-nucleoside phosphorylase, thymidylate synthase, toxicity, uridine phosphorylase

## Abstract

*Tetrahymena* are ciliated protists that have been used to study the effects of toxic chemicals, including anticancer drugs. In this study, we tested the inhibitory effects of six pyrimidine analogs (5-fluorouracil, floxuridine, 5’-deoxy-5-fluorouridine, 5-fluorouridine, gemcitabine, cytarabine) on wild-type CU428 and conditional mutant NP1 *Tetrahymena thermophila* at room temperature and the restrictive temperature (37°C) where NP1 does not form the oral apparatus. We found that cytarabine was the only tested analog that did not inhibit growth, and phagocytosis was not required for pyrimidine analog entry. IC50 values did not significantly differ between strains for the same analog at either temperature. To investigate the mechanism of inhibition, we used two pyrimidine bases (uracil and thymine) and three nucleosides (uridine, thymidine, 5-methyluridine) to help determine whether the inhibitory effects from analogs were reversible. We found that the inhibitory effects from 5-fluorouracil could be reversed by uracil and thymine, from floxuridine could be reversed by thymidine, and from 5’-deoxy-5-fluorouridine could be reversed by uracil. None of the tested nucleobases or nucleosides could reverse the inhibitory effects of gemcitabine or 5-fluorouridine. Our results suggest that the five pyrimidine analogs act on different sites to inhibit *T. thermophila* growth and that nucleobases and nucleosides are metabolized differently.

## Introduction

*Tetrahymena thermophila* are unicellular, ciliated protists whose function as a model eukaryote has led to the discovery of catalytic RNA, telomerase and telomere structure, and the first histone acetyltransferase, among numerous other contributions [1–3]. In addition to their important role in the lab, *Tetrahymena* species that live in diverse natural freshwater habitats have been used to indicate levels of pollutants present in ecosystems and to examine chemical toxicities [4,5], including different classes of anticancer drugs [4–7]. Among these anticancer drugs, we are particularly interested in pyrimidine analogs [6,8]. Their ability to block the growth of *Tetrahymena* allows us to study nucleotide uptake and metabolism in this species. Moreover, because *Tetrahymena* lacks *de novo* nucleotide synthesis pathways [9,10], we can gain further insight into the metabolism of these compounds in this organism by manipulating the external supply of nucleotides [11].

Because *Tetrahymena* cannot synthesize their own nucleotides *de novo* (review in [8]), they must obtain nucleobases, nucleosides, and nucleotides from food sources or their environment. Studies have shown that uracil, uridine, deoxyuridine, cytidine, deoxycytidine, CMP, dCMP, UMP, and dUMP could each effectively serve as the lone pyrimidine source to support the growth of *T. pyriformis* in defined medium [9,11–13]. These observations suggest that most cytidine and uridine derivatives, except for cytosine which is inert in supporting the growth of *T. pyriformis* [9,11,12], may be deaminated and aminated to form necessary pyrimidines. Moreover, comparisons of *T. pyriformis* doubling time in defined medium suggested that uracil and uridine are equally effective in supporting *Tetrahymena* growth [11].

In contrast to the effective pyrimidine sources mentioned earlier, methylated pyrimidines such as thymine, thymidine, TMP, and 5-methyluridine could not function as the lone pyrimidine source, suggesting that *T. pyriformis* lacks a pathway to demethylate these thymidine derivatives [11–13].

As a unicellular organism, how nutrients are obtained and enter *Tetrahymena* cells is also of great interest [14]. In addition to the distinct phagocytosis through their oral apparatus, *Tetrahymena* has at least four other pathways of endocytic uptake through their peripheral membrane [15]. While phagocytosis plays an important role in obtaining nutrients in *Tetrahymena*, there are also indications that some amino acids, nucleobases, and nucleosides may be absorbed through their peripheral membrane system or endocytosis [16–18]. Rasmussen first reported that at a growth condition where food vacuoles formed slowly, the doubling time of *T. pyriformis* could be shortened by supplementing the sterile filtered proteose peptone medium with high concentrations of nucleotides and glucose [18]. Using the conditional mutant NP1, a strain that does not form food vacuoles at the restrictive temperature (37°C), Rasmussen and Orias showed that at 37°C, ‘mouthless’ NP1 could multiply quickly (every 3.5 hours) in a two percent proteose peptone medium supplemented with high concentrations of vitamins and heavy metal salts [17]. Their finding further indicates that nucleobases and their derivatives, which are essential for *Tetrahymena* growth, could enter *Tetrahymena* cells through the peripheral membrane system. Freeman and Moner fed *T. pyriformis* GL-7 with [^3^H]uridine in a short period of time before a new food vacuole could be formed and quantified the amount of radioactive uridine present in different forms in the cell [16]. They concluded that uridine could enter through the cell surface and was immediately phosphorylated.

In this study, we tested the growth inhibition of six pyrimidine analogs (5-fluorouracil, floxuridine, 5’-deoxy-5-fluorouridine, 5-fluorouridine, gemcitabine, cytarabine) on the wild-type CU428 and conditional mutant NP1 *T. thermophila.* While several 5-fluorouracil derivatives have been shown to inhibit *Tetrahymena pyriformis* growth [6,11], more recently developed pyrimidine analogs, such as gemcitabine, a cytidine analog, and 5’-deoxy-5-fluorouridine, have not been evaluated in *Tetrahymena*. Furthermore, little is known about how these pyrimidine analogs are transported into and act in the cells. Our results show that except for cytarabine, the other five analogs are capable of inhibiting the growth of *T. thermophila* with different half-maximal inhibitory concentrations (IC50). We also show that phagocytosis is not required for pyrimidine analog uptake. Finally, by supplementing *T. thermophila* with higher concentrations of exogenous nucleobases and nucleosides, some but not all inhibitory effects from the analogs could be reversed in the rescue experiments. Implications from our study and how our results conform to bioinformatic predictions based on the *T. thermophila* genome are also discussed [19,20].

## Materials and Methods

### *Tetrahymena thermophila* growth curve measurements

*T. thermophila* wild-type CU428 (Stock ID: SD00178, *MPR1;* mp-s, VII) and conditional mutant NP1 strains (Stock ID: SD01422) [21,22] were acquired from the Tetrahymena Stock Center. Cells were maintained and grown at room temperature (air-conditioned at 22°C) in modified Neff’s medium (0.25% proteose peptone, 0.25% yeast extract, 0.5% glucose, 33.3 μM FeCl_3_) in Erlenmeyer flasks until the exponential growth phase. The absence of oral apparatus and phagocytosis of the NP1 strain at the restrictive temperature (37°C) was confirmed by feeding the cells Congo Red dye and examining the presence of red food vacuoles under light microscope (S1 Fig). The final cell concentration to start growth curve experiments was approximately 4,000 cells/mL.

Stock solutions of pyrimidine bases, nucleosides, and analogs were prepared by dissolving each chemical: uracil (Sigma-Aldrich, St. Louis, MO, USA), thymine (Sigma-Aldrich), uridine (Cayman Chemical, Ann Arbor, MI, USA), thymidine (Cayman Chemical), 5-methyluridine (Cayman Chemical), 5-fluorouracil (Cayman Chemical), floxuridine (Cayman Chemical), 5’-deoxy-5-fluorouridine (TCI, Portland, OR, USA), 5-fluorouridine (TCI), gemcitabine (Cayman Chemical), and cytarabine (Cayman Chemical) in modified Neff’s medium. The stock concentration for each chemical is provided in the Supplementary Materials and Methods.

Growth inhibition and rescue experiments were conducted in triplicates at room temperature for 48 hours. In rescue experiments, additional nucleobases (uracil, thymine) or nucleosides (uridine, thymidine, 5-methyluridine) were added to *T. thermophila* culture with the pyrimidine analogs in modified Neff’s medium. The final concentration of additional nucleobases and nucleosides was set at 5 mM, which was substantially higher than the concentrations of total ribonucleotides present in the modified Neff’s medium from the yeast extract (4% of the dry weight, or 0.3 mM [23]). Each inhibitory analog was tested in two different concentrations. The inhibitory concentrations of the analogs in the rescue experiments varied: 5-fluorouracil (0.4 mM), floxuridine (0.4 mM), 5’-deoxy-5-fluorouridine (2.5 mM), 5-fluorouridine (0.5 mM), and gemcitabine (0.04 mM). In addition, each analog was also tested at its IC50 concentration. Analogs that might be rescued by additional nucleobases or nucleosides were further tested in an additional concentration: 5-fluorouracil (0.05 mM), floxuridine (0.08 mM), and 5’-deoxy-5-fluorouridine (1.25 mM).

To determine whether phagocytosis is required for the entry of pyrimidine analogs, analogs were added to cell cultures four hours after cells were incubated at 37°C. Cell densities were measured 24 hours after cell inoculation (S2 Fig) using either a WPA CO8000 Cell Density Meter (Biochrom, Cambridge, UK) or a Multiskan FC microplate reader with a 594 nm filter (Thermofisher, Waltham, MA, USA).

### Calculation of IC50 values

Data analysis was performed on GraphPad Prism v9.2.0 (San Diego, CA, USA). Half-maximal inhibitory concentrations (IC50) of the tested pyrimidine analogs for CU428 and NP1 were calculated using a nonlinear regression analysis of log(inhibitor) vs. response - variable slope (four parameters). IC50 values for each analog and their standard errors are reported in Table 1. Independent-groups t-tests were performed to examine the statistical significance between IC50 concentrations of the same analog on CU428 and NP1.

**Table 1.**
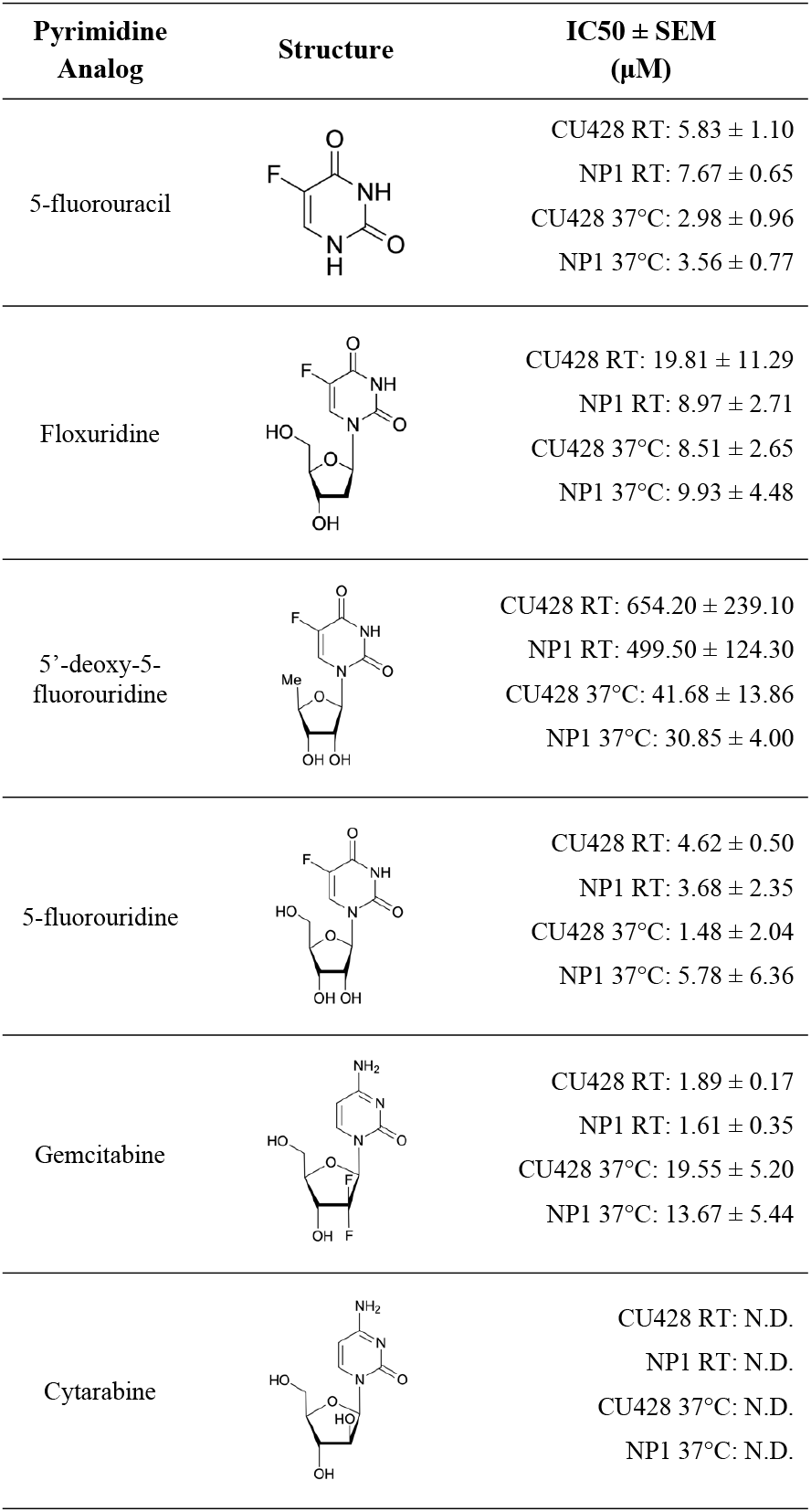
Pyrimidine analog structures and IC50 values on CU428 and NP1 *Tetrahymena thermophila* at room temperature (RT) and 37°C.

| Pyrimidine Analog | Structure | IC<sub>50</sub> ± SEM (μM) |
| --- | --- | --- |
| 5-fluorouracil |  | CU428 RT: 5.83 ± 1.10<br>NP1 RT: 7.67 ± 0.65<br>CU428 37°C: 2.98 ± 0.96<br>NP1 37°C: 3.56 ± 0.77 |
| Floxuridine |  | CU428 RT: 19.81 ± 11.29<br>NP1 RT: 8.97 ± 2.71<br>CU428 37°C: 8.51 ± 2.65<br>NP1 37°C: 9.93 ± 4.48 |
| 5'-deoxy-5-fluorouridine |  | CU428 RT: 654.20 ± 239.10<br>NP1 RT: 499.50 ± 124.30<br>CU428 37°C: 41.68 ± 13.86<br>NP1 37°C: 30.85 ± 4.00 |
| 5-fluorouridine |  | CU428 RT: 4.62 ± 0.50<br>NP1 RT: 3.68 ± 2.35<br>CU428 37°C: 1.48 ± 2.04<br>NP1 37°C: 5.78 ± 6.36 |
| Gemcitabine |  | CU428 RT: 1.89 ± 0.17<br>NP1 RT: 1.61 ± 0.35<br>CU428 37°C: 19.55 ± 5.20<br>NP1 37°C: 13.67 ± 5.44 |
| Cytarabine |  | CU428 RT: N.D.<br>NP1 RT: N.D.<br>CU428 37°C: N.D.<br>NP1 37°C: N.D. |

## Results and Discussion

### IC50 values of pyrimidine analogs on CU428 and NP1

Of the six pyrimidine analogs we tested, cytarabine (up to 0.5 mM) was the only one that did not show growth inhibition on CU428 cells (Table 1). This is consistent with results reported in *T. pyriformis,* where no growth inhibition was observed with 0.22 mM of cytarabine (arabinosylcytosine) [6]. The other five analogs inhibited CU428 and NP1 cell growth with IC50 values ranging from the low single digits to hundreds of micromolar, with the lowest being 1.61 μM (gemcitabine) and the highest being 654.20 μM (5’-deoxy-5-fluorouridine)(Table 1). Independent-groups t-tests comparing each analog’s IC50 values for CU428 and NP1 at room temperature found no significant differences for any of the five analogs (*p* > 0.3, S1 Table).

Our findings are in agreement with Hill et al.’s results, which were obtained using *T. pyriformis* grown in proteose-peptone medium [6], demonstrating that *T. thermophila* growth can be inhibited by 5-fluorouracil, 5-fluorouridine, and floxuridine (5-fluorodeoxyuridine). In addition, we have found that both gemcitabine and 5’-deoxy-5-fluorouridine are effective in inhibiting the growth of *T. thermophila.* Gemcitabine was recently shown to inhibit the growth of the marine ciliate *Euplotes vannus* [24], but has not been tested in other ciliate species.

### The uptake of pyrimidine bases and nucleosides in *T. thermophila* does not require phagocytosis

To test whether the uptake of pyrimidine analogs depends on phagocytosis, we cultured both CU428 and NP1 cells at 37°C, a temperature where the conditional mutant NP1 does not form the oral apparatus required for phagocytosis [21]. The absence of phagosomes was confirmed by using a Congo Red stain before the addition of pyrimidine analogs (S1 Fig). All five analogs inhibited cell growth of both the wild-type CU428 and the ‘mouthless’ NP1 cells at 37°C (Table 1), which strongly support the idea that the uptake of pyrimidine analogs, including nucleobases and nucleosides in *T. thermophila*, does not require an oral apparatus and phagocytosis.

Specifically how nucleobases and nucleosides are transported into *Tetrahymena* remains unknown. Several early studies in *T. pyriformis* suggested that their transports might involve the use of a nucleoside transporter system [16,25,26]. Transporter proteins in this system have the ability to move one or more types of nucleotides across membranes, with functional redundancies [16,26]. Indeed, results derived from recent bioinformatic analyses suggest that there are more than 10 equilibrative nucleoside transporter genes present in the *T. thermophila* genome [27,28], indirectly confirming those early observations. However, the functions and locations of these putative transporters, as well as the mode of transport, require further investigation.

No significant differences were found between each analog’s IC50 values for CU428 and NP1 at 37°C (all *p* > 0.2, S1 Table). Noteworthy, the IC50 values of 5-fluorouracil, 5-fluorouridine, and floxuridine on CU428 and NP1 did not change significantly between room temperature and 37°C (all *p* > 0.1). In contrast, IC50 values calculated from growth at 37°C of 5’-deoxy-5-fluorouridine decreased significantly (*p* = 0.001) while that of gemcitabine increased significantly (*p* = 0.011), when comparing to those values measured at room temperature (Table 1). We do not have a simple explanation for the changes (or lack thereof) in IC50 values, as the exact inhibition mechanisms for each of the analogs in *T. thermophila* are not known. An analog may act on one or more enzymes, and may have varying binding affinities to one enzyme at different temperatures.

### Growth inhibitory effects may be reversed by adding additional nucleobases or nucleosides

To further investigate how the five pyrimidine analogs might affect pyrimidine metabolism in *T. thermophila*, we added additional pyrimidine bases and nucleosides to determine whether they could compete against the analogs and reverse the growth inhibition. Two pyrimidine bases (uracil and thymine) and three nucleosides (uridine, thymidine, and 5-methyluridine) were used in the rescue experiments; the nucleobases and nucleosides were added alongside the pyrimidine analogs to *T. thermophila* culture. We expected that higher concentrations of uracil and uridine might outcompete 5-fluorouracil and its derivatives, but not gemcitabine, which is a cytidine analog. Furthermore, if thymidylate synthase, which synthesizes the essential thymidine for DNA replication by converting dUMP to dTMP, is the primary inhibitory site for 5-fluorouracil and its derivatives as shown in human cancer cells [29,30], supplying *T. thermophila* with additional thymine or thymidine should help release the inhibitory effect. We did not expect 5-methyluridine to reverse the inhibitory effects since it is inert in supporting the growth of *T. pyriformis* [11–13].

In the rescue experiments, the concentration of the five rescuing pyrimidine bases and nucleosides was fixed at 5 mM. This concentration was substantially higher than the two concentrations of each pyrimidine analog tested: an inhibitory one and the IC50 (See Materials and Methods). For 5-fluorouracil, floxuridine, and 5’-deoxy-5-fluorouridine, whose inhibitory effects were found to be reversible in the rescuing experiments, we tested an intermediate concentration between the inhibitory concentration and IC50.

As predicted, 5-methyluridine was not able to reverse inhibitory effects from any of the five analogs (Figs 1-4) and none of the five pyrimidine bases and nucleosides we tested could reverse the inhibitory effect of gemcitabine (Figs 4A-B). Additionally, the growth inhibition from 5-fluorouridine could not be reversed by any of the five pyrimidine bases and nucleosides (Figs 4C-D). In the absence of analogs, none of the additional pyrimidine bases or nucleosides affected *T. thermophila* growth (Figs 1D, 2D, 3D).

**Fig 1.**
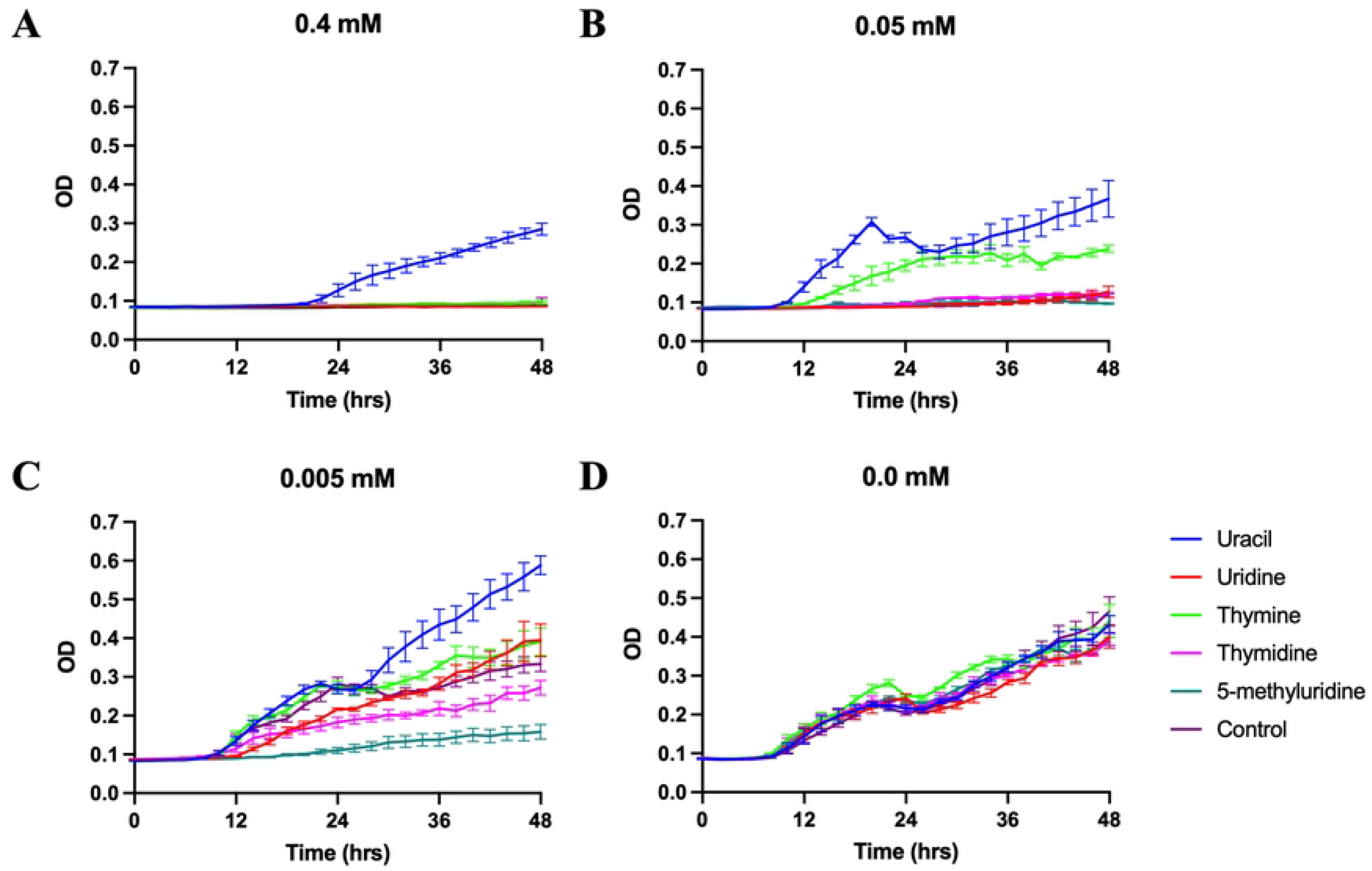
Growth curves of *T. thermophila* supplemented with 5 mM of uracil, uridine, thymine, thymidine, or 5-methyluridine in modified Neff’s medium containing: (A) 0.4 mM. (B) 0.05 mM. (C) 0.005 mM. (D) 0.0 mM 5-fluorouracil. Data plotted ± SEM.

**Fig 2.**
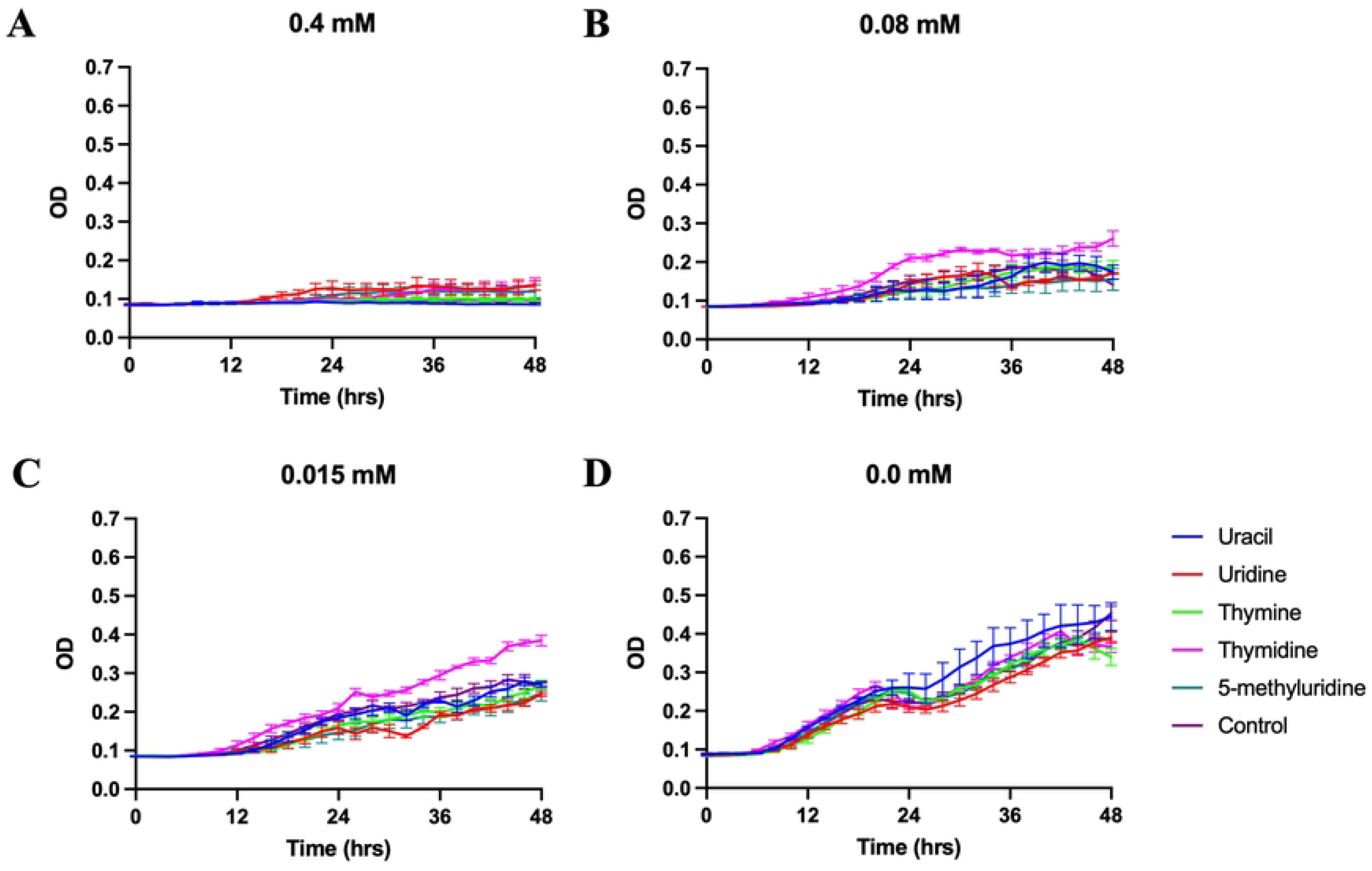
Growth curves of *T. thermophila* supplemented with 5 mM of uracil, uridine, thymine, thymidine, or 5-methyluridine in modified Neff’s medium containing: (A) 0.4 mM. (B) 0.08 mM. (C) 0.015 mM. (D) 0.0 mM floxuridine. Data plotted ± SEM.

**Fig 3.**
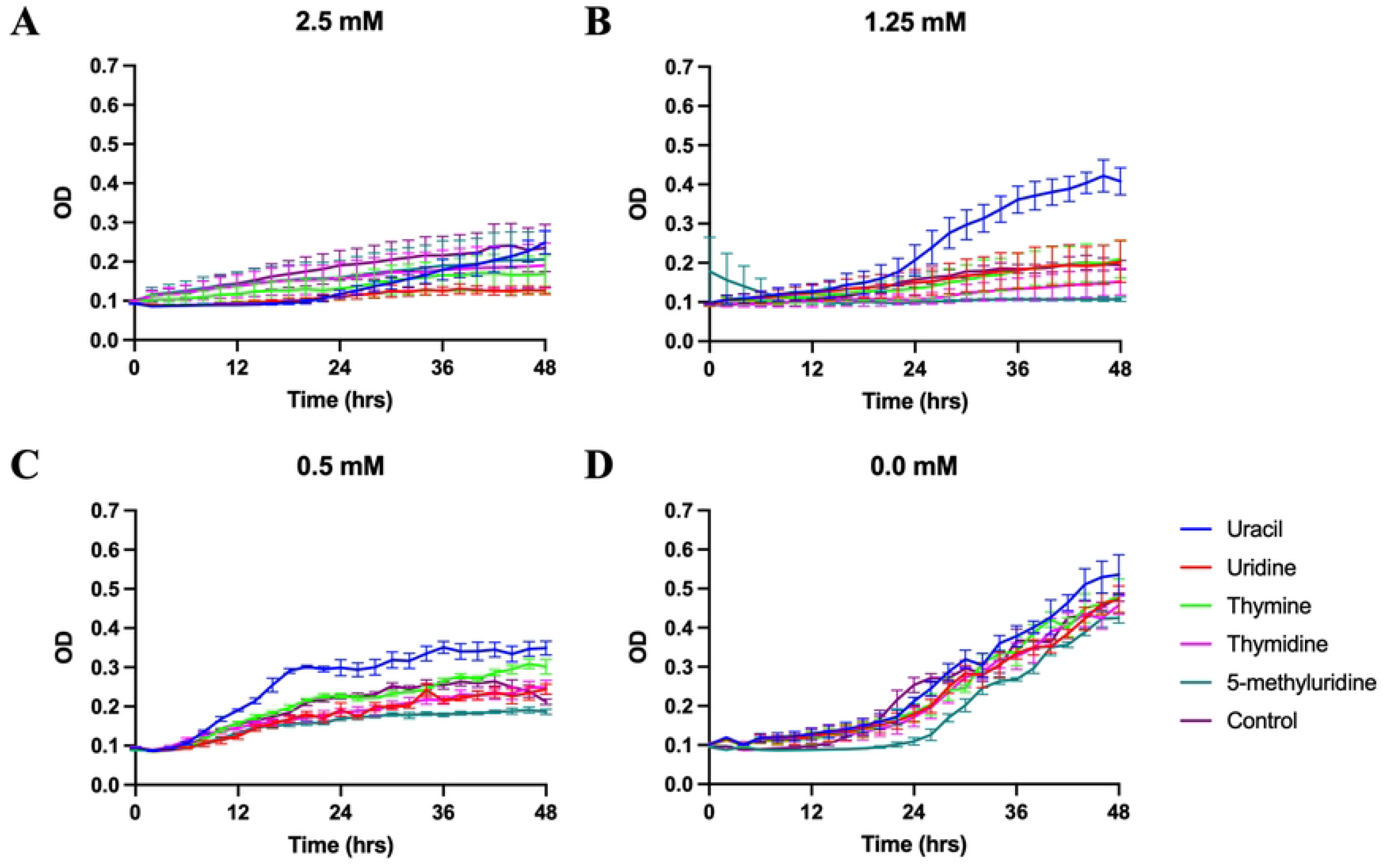
Growth curves of *T. thermophila* supplemented with 5 mM of uracil, uridine, thymine, thymidine, or 5-methyluridine in modified Neff’s medium containing: (A) 2.5 mM. (B) 1.25 mM. (C) 0.5 mM. (D) 0.0 mM 5’-deoxy-5-fluorouridine. Data plotted ± SEM.

**Fig 4.**
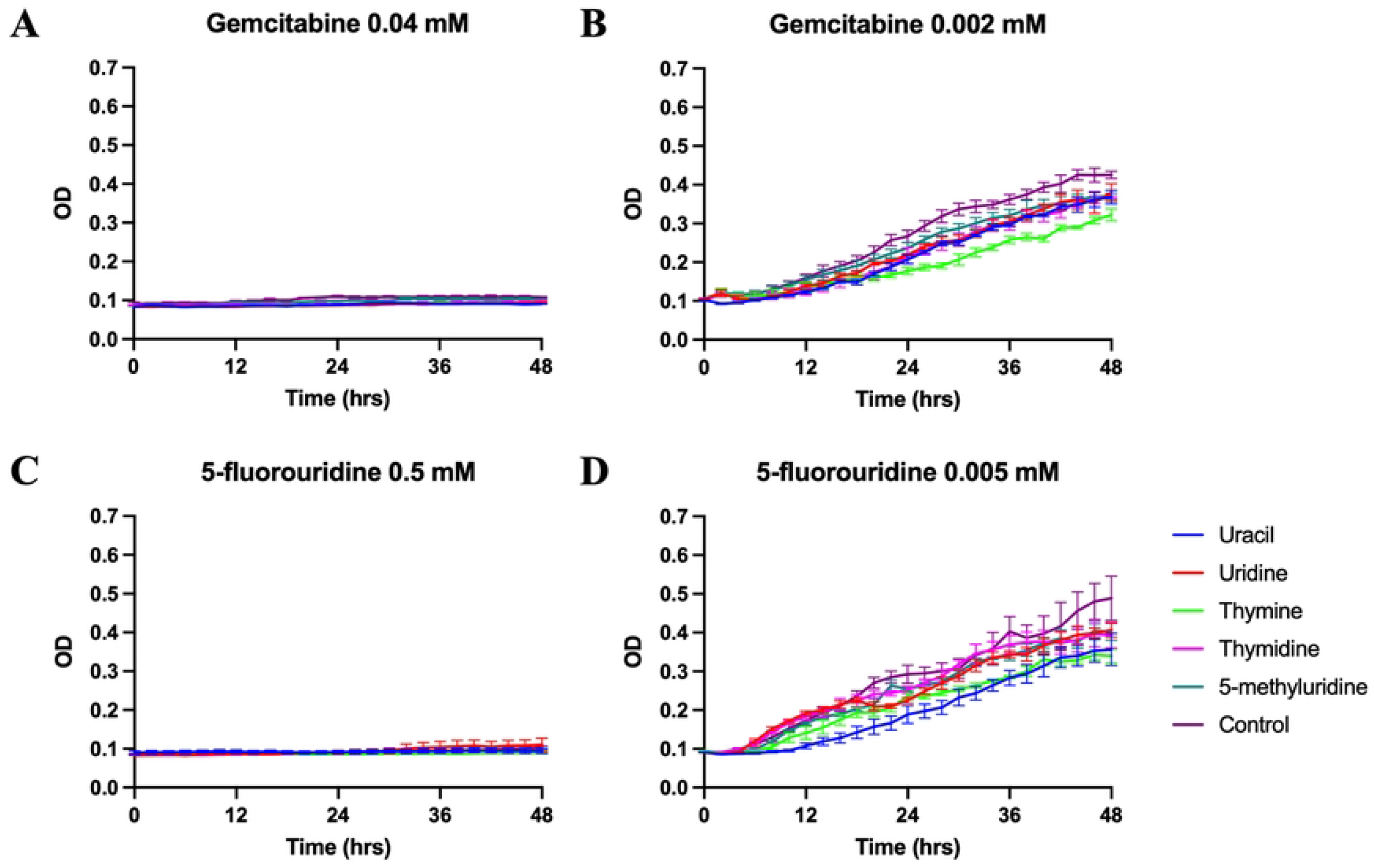
Growth curves of *T. thermophila* supplemented with 5 mM of uracil, uridine, thymine, thymidine, or 5-methyluridine in modified Neff’s medium containing: (A) 0.04 mM gemcitabine. (B) 0.002 mM gemcitabine. (C) 0.5 mM 5-fluorouridine. (D) 0.005 mM 5-fluorouridine. Data plotted ± SEM.

We found that uracil but not uridine, and thymine but not thymidine, could reverse the inhibitory effects from 5-fluorouracil (Fig 1). Uracil was capable of reversing the inhibitory effect at all three 5-fluorouracil concentrations (Figs 1A-C). The rescue effect of thymine was most profound at the intermediate concentration of 5-fluorouracil (0.05 mM, Fig 1B), but not at a higher, inhibitory concentration (0.4 mM, Fig 1A).

Growth inhibition from floxuridine could only be reversed by thymidine at the intermediate concentration and IC50 (Fig 2), and those from 5’-deoxy-5-fluorouridine could only be reversed by uracil at the two lower concentrations (Fig 3).

While early studies suggested that uracil and uridine were equally effective as the sole pyrimidine source supporting *Tetrahymena* growth [11], our rescue experiments showed that only uracil was capable of reversing the growth blockages in *T. thermophila* caused by 5-fluorouracil and its prodrug 5’-deoxy-5-fluorouridine (Figs 1 and 3). Clearly, uracil and uridine are not as interchangeable in *Tetrahymena* as once thought. One clue for this non-interchangeability may be found in the *Tetrahymena* genome [20] as the key enzyme responsible for interconverting uridine and uracil is absent, according to the KEGG pathway (S3 Fig) [19]. However, results derived from several early experiments strongly suggest the presence of uridine phosphorylase activity in *T. pyriformis* [31–33]. Whether these discrepancies indicate the presence of a non-canonical uridine phosphorylase in *Tetrahymena* remains to be further investigated. Our results suggest that uridine and uracil are differentially metabolized in *T. thermophila*: uridine may be preferentially used in RNA synthesis and uracil for DNA synthesis when both are available. Because 5-fluorouracil and 5’-deoxy-5-fluorouridine primarily target thymidylate synthase for dTMP synthesis (EC 2.1.1.45 in S3 Fig, review in [34]), uracil may be more effective in reversing their inhibitive effect.

Interestingly, in addition to uracil and uridine, thymine and thymidine also appear to be differentially metabolized in *Tetrahymena*. This was first indicated in an early experiment where thymidine, but not thymine, was shown to reinitiate growth of *T. pyriformis* grown in defined medium lacking folic acid [11]. We show that thymine, but not thymidine, reversed the growth blockage from 5’-deoxy-5-fluorouridine in *T. thermophila* grown in modified Neff’s medium (Fig 3). Thymidine, but not thymine, could similarly reverse the growth blockage from floxuridine (Fig 2). Altogether, these observations strongly suggest inefficient interconversions between thymine and thymidine, despite the biochemical and bioinformatic detection of thymidine phosphorylase activity in *Tetrahymena* [35] (S3 Fig, EC 2.4.2.4). Indeed, it has been shown that thymidine is a much more effective precursor than thymine to be incorporated into DNA in *Tetrahymena* among other eukaryotic cells [36,37]. Finally, the fact that thymine (but not thymidine) could reverse the growth blockage from 5’-deoxy-5-fluorouridine indicates that this nucleobase might be able to be converted to TMP without going through a thymidine intermediate. However, other mechanisms (e.g. competitive and allosteric regulations) to interpret our results remain equally plausible.

It is noteworthy that our results show that thymidine, but not uridine, rescued the growth of *T. thermophila* from floxuridine blockage in modified Neff’s medium (Fig 2). This would contradict the results from *T. pyriformis* grown in a defined medium that uridine, but not thymidine, reversed the growth blockage from the same analog [11]. It is not clear what may have caused the discrepancies. The concentrations of floxuridine used in both studies were similar while our experiments used a much higher concentration of rescuing pyrimidines (5 mM) compared to approximately 41 μM (10 mg/ml) in Wykes and Prescott’s study [11]. In addition, the *Tetrahymena* species and growth media used in the two studies were different. More experiments are clearly needed to further elucidate the blockage mechanisms of floxuridine and other pyrimidine analogs and the mechanism of nucleotide transportation and metabolism in different *Tetrahymena* species.

## Conclusions

In this study, we demonstrated that 5-fluorouracil, floxuridine, and 5-fluorouridine inhibited the growth of *T. thermophila* grown in modified Neff medium (compared to *T. pyriformis* in a defined medium in previous studies). Furthermore, we showed that both gemcitabine and 5’-deoxy-5-fluorouridine inhibit *T. thermophila* growth, while cytarabine does not. We present new evidence suggesting that these pyrimidine analogs may enter *T. thermophila* through their peripheral membrane systems. Results obtained from the rescue experiments suggest differential metabolism of uracil and uridine, and thymine and thymidine in *T. thermophila*.

## Acknowledgements

The authors would like to thank Drs. Thomas Doak, F. Paul Doerder, Donna Cassidy-Hanley, and Ed Orias for providing background information and guidance on *Tetrahymena* strains. JC and SW were supported by Hamilton Summer Research Funds.

## Author Contributions

Conceptualization, WJC and ZW; methodology, data acquisition and analyses, ZH, WJC, JC, RC, IC, JR, and SW; manuscript preparation, ZH, WJC, and ZW; supervision, WJC and ZW. All authors have read and agreed to the published version of the manuscript.

## Supporting Information

**S1 Fig. Micrographs of CU428 and NP1 *T. thermophila* fed Congo Red dye at RT and 37°C. Micrographs taken at 20X magnification, 1.5X zoom. (A) CU428 at RT. (B) NP1 at RT. (C) 428 at 37°C. (D) NP1 at 37°C. The lack of dark vacuoles in NP1 at 37°C indicates a nonfunctional oral apparatus.**

**S2 Fig. *T. thermophila* growth (O.D.) as a function of increasing pyrimidine analog concentrations. Data plotted ± SEM.**

**S3 Fig. KEGG pyrimidine metabolism pathway of *T. thermophila.* Homologous genes detected in the *T. thermophila* genome are highlighted in green. This diagram was created and downloaded from the KEGG database** [19] **by selecting the pathway type for *T. thermophila*.**

**S1 Table. Comparisons of IC50 values of pyrimidine analogs on CU428 and NP1 at room temperature and 37°C.**

